# Interferons Reshape the 3D Conformation and Accessibility of Macrophage Chromatin

**DOI:** 10.1101/2021.03.09.434568

**Authors:** Ekaterini Platanitis, Stephan Grüner, Aarathy Ravi Sundar Jose Geetha, Laura Boccuni, Alexander Vogt, Maria Novachkova, Andreas Sommer, Mathias Müller, Thomas Decker

## Abstract

Engagement of macrophages in innate immune responses is directed and enhanced by type I and type II interferons. An essential component of IFN activity is the use of JAK-STAT signal transduction for the transcriptional control of interferon-stimulated genes (ISG). Here, we study the immediate early nuclear response to type I IFN and IFN-γ in murine macrophages. Despite their distinct immunological activities, both IFN types triggered highly overlapping epigenomic and transcriptional changes. These changes included a rapid rearrangement of the 3D chromatin organization and an increase of DNA accessibility at ISG loci. ISGF3, the major transcriptional regulator of ISG, controlled homeostatic as well as induced-state DNA accessibility at a subset of ISG. Increases in DNA accessibility correlated with the appearance of activating histone marks at surrounding nucleosomes. Collectively our data emphasize changes in the three-dimensional nuclear space and epigenome as an important facet of transcriptional control by the IFN-induced JAK-STAT pathway.

## Introduction

Interferons are soluble messengers that are produced in response to invading pathogens. Particularly the type I-IFN species IFN-α and IFN-β (collectively called IFN-I) are of vital importance in the course of cell autonomous antiviral immunity. IFN-γ, the only member of the type II interferon family, is similarly capable of inducing the antiviral state, but functions predominantly as a macrophage-activating cytokine in the immune system^1–3^. Although IFN-I and IFN-γ have diverging functions in many biological scenarios, both activate the JAK-STAT pathway ^4,5^. The extent to which the two cytokines elicit different transcriptional profiles in cells is incompletely understood. Improved knowledge about nuclear responses to IFN-I and IFN-γ is required to decipher the relationship between transcriptome changes and the cytokines’ immunological impact. Additionally, such insight will improve our understanding of the molecular hallmarks of interferon-associated diseases ^6,7^.

The JAK-STAT pathway represents a striking example of receptor-mediated signal transduction ^8^. This evolutionarily conserved pathway utilizes a simple membrane-to-nucleus mechanism for rapidly inducing gene expression ^9^. Thus, it serves as a simple paradigm for how cells sense extracellular signals and translate them to a specific transcriptional outcome.

Signal transduction downstream of IFN-I receptors requires activated JAKs to cause the formation of the transcription factor IFN-stimulated gene factor 3 (ISGF3). This heterotrimer of STAT1, STAT2 and interferon regulatory factor 9 (IRF9), assembles at interferon-stimulated response elements (ISREs), present in the promoters of interferon-stimulated genes (ISG) ^10–12^. The IFN-γ receptor on the other hand employs JAKs to activate the gamma interferon-activated factor (GAF), a STAT1 homodimer, which stimulates ISG expression by binding to gamma interferon-activated sites (GAS) ^13^.

Regulation of transcription in eukaryotic cells is based on a fine-tuned interplay between chromatin structure and recruitment of a plethora of transcription factors to enhancers and promoters ^14,15^. These regulatory elements differ in their function and location with regard to the transcription start site (TSS), but share some common characteristics such as specific active epigenetic marks and their property to recruit the RNA polymerase to the core promoter and thus initiate transcription. However, putative enhancers cannot be predicted solely based on genomic proximity since gene expression programs are orchestrated within the three-dimensional nuclear space, thus often bridging considerable genomic distances ^16,17^.

Recently, genome-wide chromosome conformation capture (Hi-C) experiments have shown that chromosomes acquire their spatial order at several different levels of their structural organization ^18^. This three-dimensional folding of the eukaryotic genome is a highly organized process, tightly linked to transcription. At the megabase scale, chromosomes are organized into functionally distinct A and B compartments, representing transcriptionally active and inactive genomic regions, respectively ^19,20^. At the next level, chromosomes fold into topologically associating domains (TADs) and chromatin loops. Their formation depends on the ring-shaped, multiprotein cohesin complex that cooperates with the sequence-specific DNA binding protein CTCF ^21–24^. TADs and loops contribute to the regulation of gene expression by facilitating the interactions between promoters and enhancers located within the same domain ^25–27^.

Whether IFN induce changes of the genome organization in order to control gene expression is largely unexplored. Here, we decipher genome-wide chromatin dynamics in primary murine macrophages exposed to IFN-I and IFN-γ and find that they trigger a rapid rearrangement of the 3D chromatin organization on ISG-rich regions. Surprisingly, both interferon types cause highly overlapping transcriptional changes and chromatin rearrangement profiles, highlighting our previously reported importance of the ISGF3 complex in the IFN-γ response ^28,29^. We further observe a significant gain in genome-wide chromatin accessibility at the TSSs of ISG in response to both stimuli, which strongly correlated with mRNA expression. An active promoter configuration is further underlined by the gain of active histone marks (H3K27ac) and the removal, or lack of deposition, of transcriptional silencing modifications (H3K27me3). Finally, we present a new model, which highlights the importance of the ISGF3 complex in priming the homeostatic chromatin landscape on ISG, as well as in remodeling the local chromatin structure downstream of both type I and type II IFN signaling.

## Results

### IFN increase the A compartment strength at ISG

To examine and compare the immediate early response effects of IFN treatment on macrophage chromatin 3D conformation on a whole-genome level, we treated mouse bone marrow-derived macrophages (BMDM) for 2 hours with either IFN-I or IFN-γ and analyzed chromatin contacts by Hi-C in two biological replicates.

First, we determined the reproducibility of our data with HiCRep, a framework using a stratum-adjusted correlation coefficient for quantifying sample similarity from Hi-C interaction matrices ^30^ (Fig. EV 1A). The replicates clustered together according to their treatment, allowing us to merge replicates and thus obtain a higher coverage for subsequent analysis. In addition to the overall sample similarity, the data show that both IFN treatments produce changes of the overall chromatin structure.

Genome-wide analysis studies have revealed that euchromatin and heterochromatin regions are spatially segregated ^19,21,27^. Thus, Hi-C pair-wise interaction heatmaps not only show that individual active and inactive regions remain spatially separated, but that each type of chromatin also interacts with distal regions of the same type. Here we used eigenvector deconvolution analysis of our Hi-C data to identify regions that correspond to each type of chromatin. ‘A’ compartments represent active regions that are usually defined as gene-rich with ongoing transcription and high accessibility. ‘B’ compartments on the other hand are inactivated regions containing tightly packed and inaccessible chromatin. Integrating our previously published RNA-seq data sets with Hi-C experiments performed under matching experimental conditions allowed us to confirm the accuracy of our compartment analysis by plotting the compartment scores of each genomic bin against the number of RNA-seq reads mapping to the respective bin (Fig. EV1 B-D) ^28^. We observed a correlation between the compartment scores and RNA-seq read enrichment that underlines the segregation of the genome into A and B compartments based on transcriptional activity. Visualization of contact maps in combination with principal component analysis, revealed that the strongest effects within compartments occur in clustered ISG (Fig. 1A, C). Active compartments are represented by a positive eigenvector and displayed in red, while B compartments are shown in blue. Here we highlight the *Gbp* gene cluster (depicted by the black bar), which showed rearrangements in the contact map as well as an increase in compartment scores and thus compartment strength upon IFN-I and IFN-γ treatment (PC1 tracks). Additionally, we observed differences in contacts between the *Gbp* cluster and surrounding regions (Fig. 1B, D). Specifically, the plots show that IFN treatment corresponds with a decrease in contacts between the *Gbp* cluster and neighboring regions. The region encompassing the *Gbp* cluster is delineated with black lines. Decreases in long-range contact probability are represented by greater intensity of blue color, increases in short-range contact probability by intensification of red color.

**Fig. 1:**
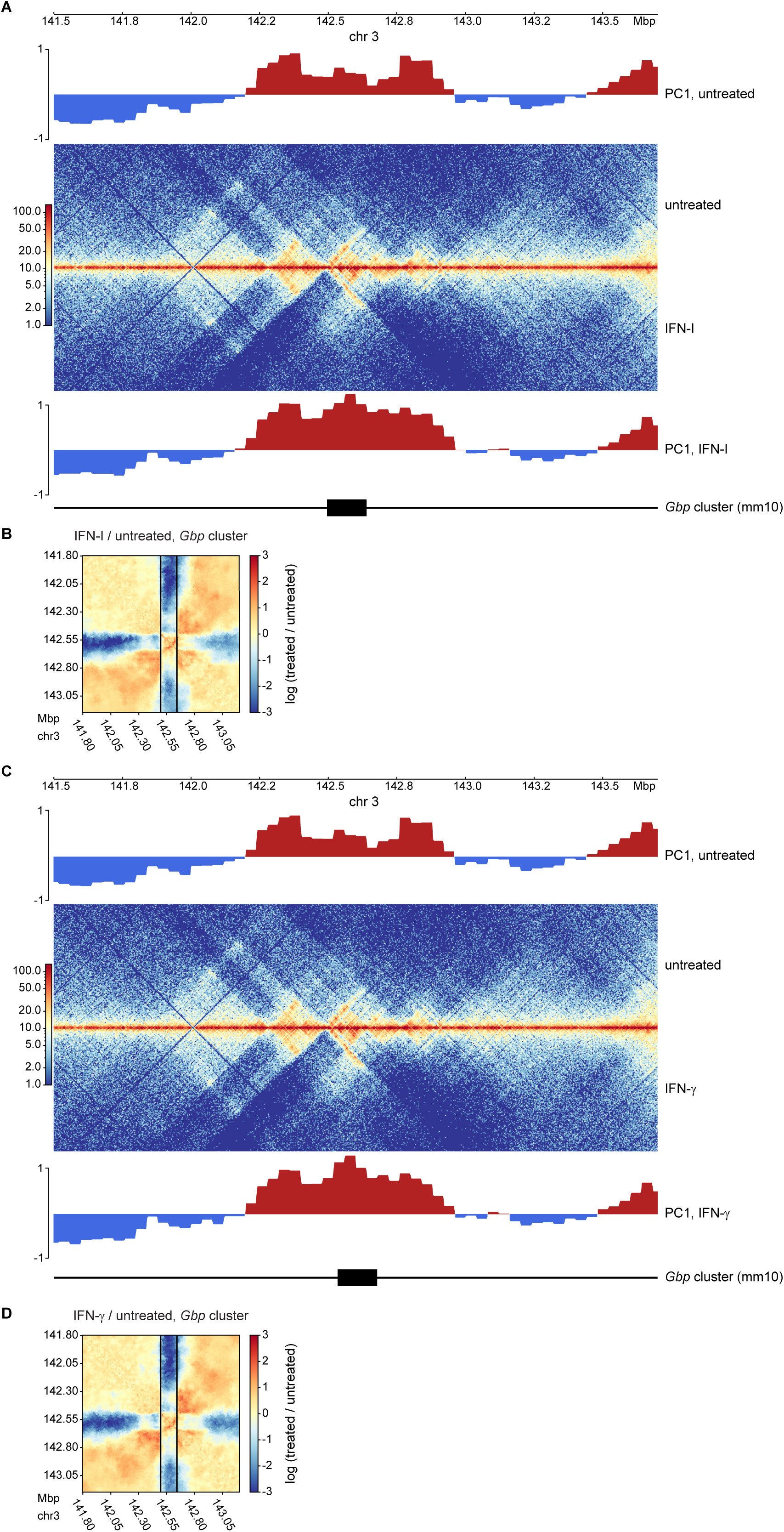
IFN treatment rearranges host cell genome 3D structure. **A**, C Hi-C contact maps around the *Gbp* locus of untreated (upper middle) and **A** IFN-I treated (2h) or **C** IFN-γ treated (2h) (lower middle) primary murine bone marrow-derived macrophages (BMDM; merge of 2 replicates per condition). Upper and lower panel, PC1 values indicating compartments. **B, D** Ratio plots around the *Gbp* locus comparing contact maps of **B** IFN-I or **d** IFN-γ treated and untreated BMDM, generated using the flexible binning method Serpentine. Black vertical lines indicate the borders of the *Gbp* locus. Red regions indicate increased, blue regions decreased interaction after treatment with IFN.

The lack of compartment switching demonstrates that the observed regions are already in an active state before IFN treatment. This observation is in line with previous findings showing that many ISG are already transcribed at low, basal levels in the absence of any stimulus ^28,31^. Yet, the change in compartment strength upon IFN-I and IFN-γ treatment correlates with a higher transcriptional activity of the genes located within those compartments. Changes in compartmentalization strength can be further visualized and quantified by plotting interaction frequencies between pairs of 80 kb bins arranged by their values along the first eigenvector to obtain saddle plots ^22^. Here, the lower right quadrant represents A-A interactions and the upper left corner represents B-B interactions (Fig. EV1E-G). We found that on a genome-wide scale the interactions of A/B compartments are largely unaffected by short term IFN-I and IFN-γ signaling. This result is in line with previous studies showing that short term exposure to other extracellular signals likewise has no effect on higher-order chromatin structures ^32–34^.

Taken together, we conclude that both IFN types, while reinforcing A compartments at ISG loci, do not globally change the genomic arrangement of compartments. This suggests that 3D alterations may result from changes in DNA loops.

### IFN-I signaling elicits rapid rearrangement of chromatin-chromatin interactions in ISG rich regions

To further compare the effects of each IFN type on the chromatin structure, we systematically analyzed differences in the contact maps and performed loop calling using the pattern detection algorithm chromosight ^35^. Here, each detected loop is represented by its start and end position, as well as a Pearson correlation score. For visualization, the detected chromatin loops are represented as arcs, whose darkness of color increases proportionally to the loop scores. We confirmed that single replicates show similar results in regard to the loop calling process, allowing us to merge the replicates for obtaining denser contact maps (Fig. EV2).

A large fraction of ISG is organized in clusters on different chromosomes. Thus, we examined whether chromatin contacts within such clusters are subject to IFN-induced changes by exploring the genomes of resting or IFN-I treated BMDM. Indeed, the resulting contact maps, representing a depth of about 500 Mio read pairs/contacts, revealed clear changes at all examined ISG clusters (Fig. 2A-F). In regions where clear changes in the contact maps and compartment scores occurred, loop positions and scores were also affected. Alterations in these clusters represented predominantly increases in intra-cluster, short-range chromatin contacts. This effect was especially noticeable at the *Gbp, Ifit* and *Oas* gene clusters (Fig. 2A-C).

**Fig. 2:**
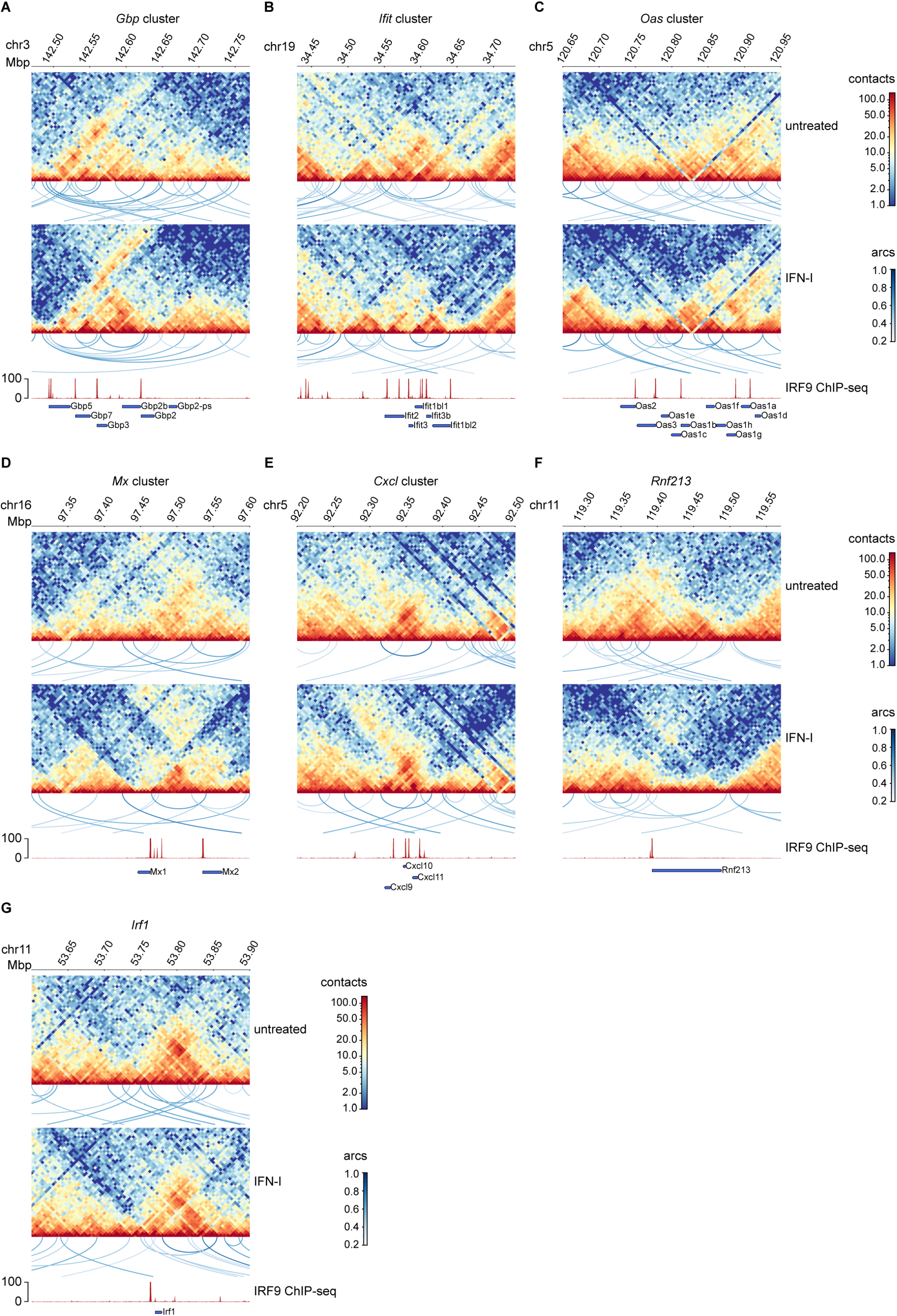
IFN-I treatment reshapes chromatin-chromatin contacts in ISG clusters. **A-G** Hi-C contact maps (merge of 2 replicates per condition, log1p scale) of untreated and IFN-I treated (2h) BMDM. Visualization of pattern detection with chromosight (loops between two loci are indicated as arcs). Color coding of arcs corresponds to Pearson correlation scores. Loops with a score > 0.35 are shown (maximum size/ylim = 500 kbp). Lower panel, IRF9 ChIP-seq tracks after IFN-I (1.5h) stimulation and gene annotations are shown to visualize ISGF3 binding sites. Only ISG of the corresponding gene clusters are visualized.

Within the *Gbp* cluster on chromosome 3, short range contacts between loci inside the region of interest showed an increased interaction after IFN-I treatment. Additionally, many long-range contacts decreased in strength after treatment, consistent with the analysis presented in Fig. 1B. Striking effects of IFN-I stimulation were also observed close to the end of chromosome 16, where *Mx1* and *Mx2* are located. Here, we found that treatment of the cells dramatically increased contacts in the area between the two genes (Fig. 2D). *Rnf213*, which is not embedded within a cluster, shows that short-range conformation changes even occur on single ISG (Fig. 2F).

Taken together, these results demonstrate that chromatin rearrangements in response to IFN-I treatment can mostly, but not exclusively, be observed around regions containing larger numbers of interferon stimulated genes and some of the clusters show similar patterns characterized by a predominant loss of long-range contacts.

### IFN-I and IFN-γ show a high overlap of transcriptional profiles and 3D chromatin rearrangements

When comparing IFN-I to IFN-γ treatment, we realized that both cytokines triggered very similar 3D chromatin changes at the majority of ISG clusters which are characterized by a prevalence of ISRE sequences (Fig. 3A, Fig. EV 3A-C). In contrast, *Irf1*, an ISG carrying only a GAS element, and thus a STAT1 binding site within its proximal promoter ^36,37^, showed stronger changes in its contact map when cells were treated with IFN-γ (Fig. 3B). A similar trend was noted for the *Irf8* gene which is similarly regulated by STAT1 homodimers ^38,39^. In line with the much weaker inducibility of the *Irf8* gene compared to *Irf1* (Fig. 3C, table EV1), IFN-γ-induced changes in its loop structure were comparably small. Although evidence that STAT binding initiates loop formation is currently elusive, the behavior of the *Irf1* and *Irf8* genes is in agreement with the paradigm of IFN signaling, which posits a larger contribution of STAT1 homodimers to IFN-γ-induced transcription when compared to those produced by IFN-I. On the other hand, loop formation at ISG clusters is in line with our previous notion that ISGF3 activity carries significant weight in the entirety of IFN-γ-induced transcriptome changes. To further address differences and similarities, we directly compared the early transcriptional response to both IFN types in BMDM based on RNA-seq datasets previously generated in our lab. Surprisingly, many genes and especially the strongly induced ones were affected in a similar fashion by IFN-I and IFN-γ (Fig. 3C). However, groups of lesser induced genes showed preferences for IFN-I (green dots) or IFN-γ (purple dots). Overall, this finding is in agreement with similar chromatin rearrangements produced by both IFN types at many ISG-rich regions. At the same time, it supports the previously reported role of the ISGF3 complex in the IFN-γ response ^28,29,40^.

**Fig. 3:**
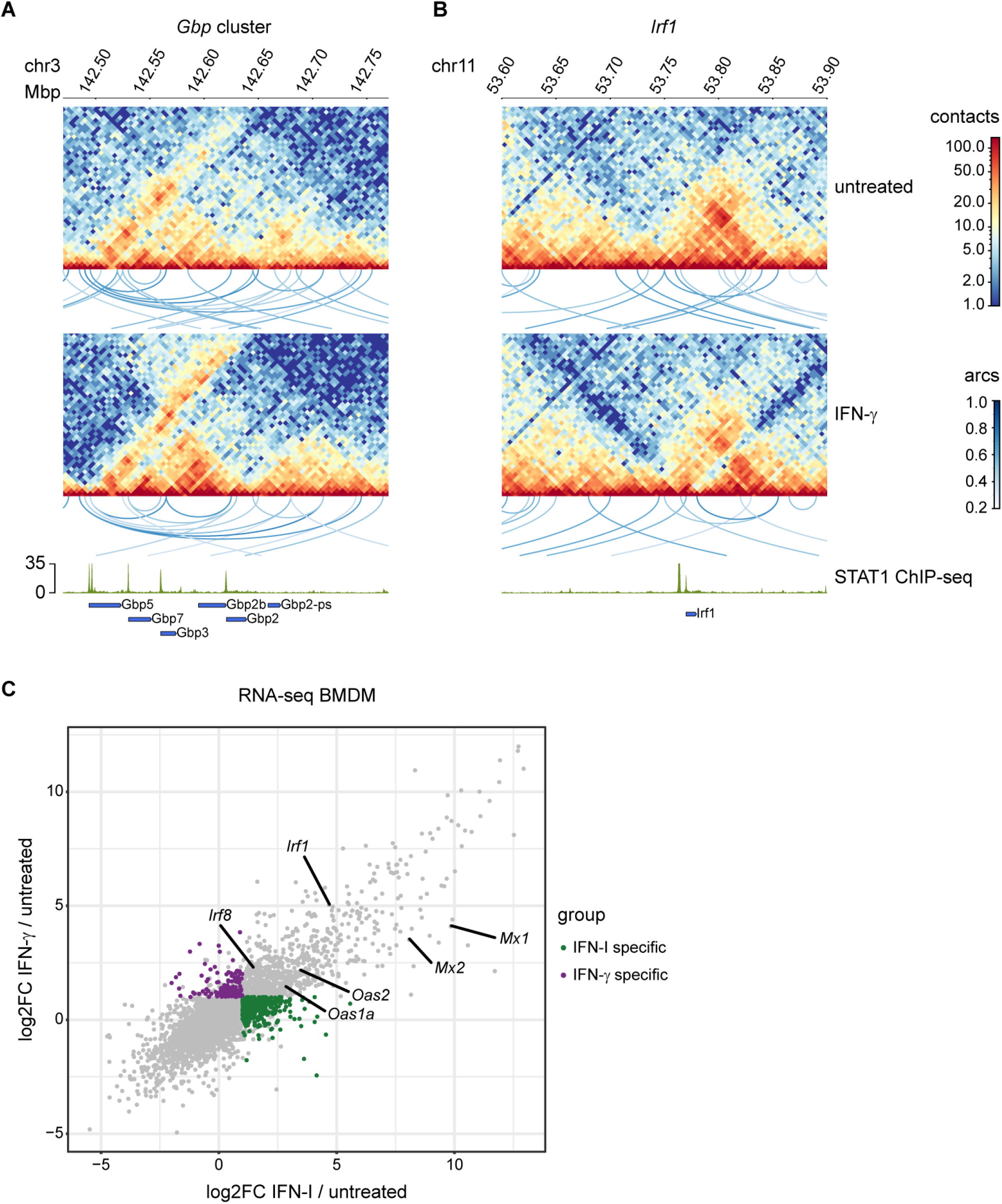
Effects of IFN-I and IFN-γ treatment on chromatin organization in ISG clusters are largely overlapping. **A**, **B** Hi-C contact maps (merge of 2 replicates per condition, log1p scale) of untreated and IFN-γ (2h) treated BMDM. Visualization of pattern detection with chromosight (loops between two loci are indicated as arcs). Color coding of arcs corresponds to Pearson correlation scores. Loops with a score > 0.35 are shown (maximum size/ylim = 500 kbp). Lower panel, STAT1 ChIP-seq tracks (1.5h IFN-I treatment) and gene annotations are shown to visualize binding sites for both ISGF3 and STAT1 homodimers. Only ISG of the corresponding gene clusters are visualized. **C** log2FC plot comparing effects of IFN-I (2h) and IFN-γ (2h) treatment in BMDM. mRNA expression ratios, with a cutoff padj ≤ 0.05 are plotted. Genes significantly upregulated (log2FC ≥ 1, padj ≤ 0.05) in one, but not the other treatment (log2FC ≤ 1) are colored in green (IFN-I) and purple (IFN-γ).

### ISGF3 regulates chromatin accessibility at ISG loci in homeostatic and IFN-induced conditions

A hallmark of the immune system is its ability to quickly transition between resting and stimulated states. Recent studies have shown large-scale chromatin remodeling in cells responding to immune stimuli ^41,42^. In order to complement our Hi-C data with information about chromatin accessibility, we performed ATAC-seq, designing experiments to reveal the impact of both IFN types on nucleosome structure. All ATAC-seq samples were generated in three biological replicates and a high correlation between replicates was obtained (Fig. EV4A).

For the analyses illustrated in Fig 4 and Fig. EV 4, the genome was divided into the top 200 IFN-I or IFN-γ -induced genes and the remaining genome (non-ISG), according to our RNA-seq analysis ^28^ (Fig. 4A-D; Fig. EV 4B,C). In light of our recent data ^28^ showing a role of ISGF3 complexes in both homeostatic and induced ISG expression, we examined *Irf9-/-* macrophages. These cells reveal the impact of ISGF3. Of note, the acronym ISGF3 describes two related complexes: the full version formed by STAT1, STAT2 and IRF9 dominating the IFN response, as well as the ‘light’ version formed by STAT2 and IRF9, whose main importance lies in the activation of homeostatic ISG transcription ^28^.

**Fig. 4:**
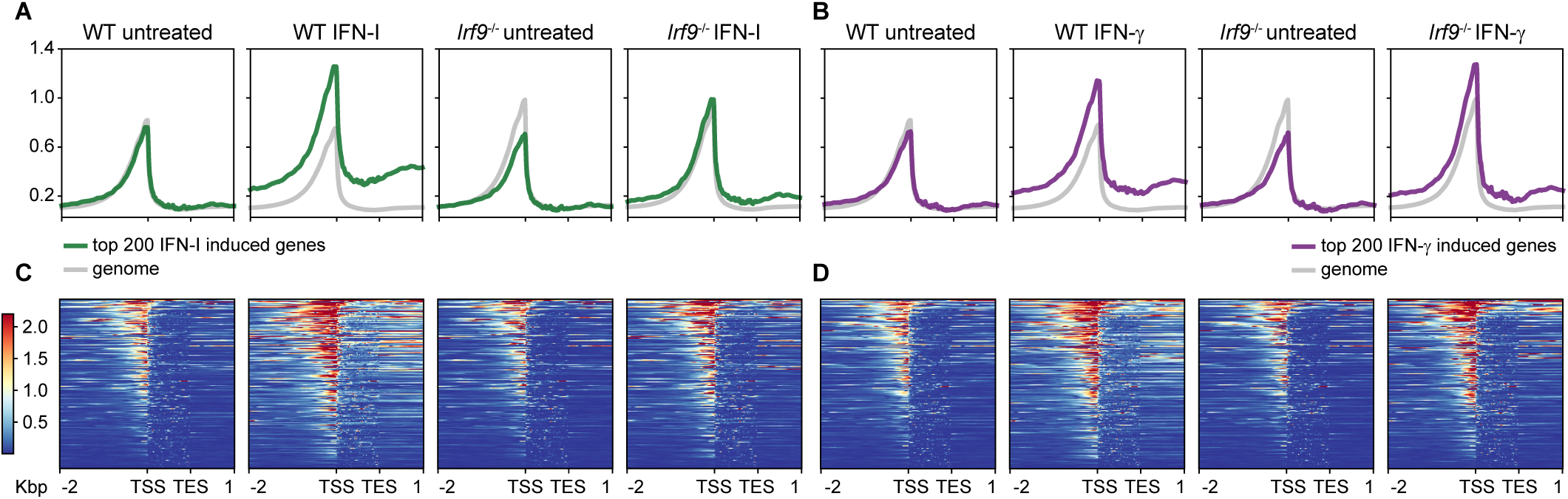
ISGF3 complex-dependent effects of IFN signaling on chromatin accessibility. **A**, **B** Summary profile plots of chromatin accessibility in BMDM generated by using normalized read coverages for untreated and IFN-I/IFN-γ treated wildtype and *Irf9-/-* BMDM as indicated. The colored profile is an average of the top 200 genes (log2FC) upregulated after IFN-I (2h; green profile) and IFN-γ (2h; purple profile) treatment respectively, according to RNA-seq. The grey profile is an average of the respective remaining gene regions. **C, D** Heatmaps of chromatin accessibility in regions of the top 200 genes (log2FC) upregulated after IFN-I (2h) and IFN-γ (2h) treatment respectively according to RNA-seq. Profiles and heatmaps are shown for the regions of the genes, including 2 kbp upstream of the TSS and 1kbp downstream of the TES as indicated below the plots. Shown are merged data from three biological replicates.

Chromatin at non-ISG did not undergo major changes upon treatment with either IFN type (Fig. 4A, B, grey profile; Fig. EV4B, C). The number of ATAC-seq signals at non-ISG was consistently higher in IRF9-deficient cells, possibly showing a role of IRF9 in repressing accessibility of certain non-ISG (Fig. 4A, B, grey profile). Importantly, the loss of IRF9 produced an accessibility decrease in a subset of ISG loci during cell homeostasis, which is consistent with the role of STAT2/IRF9 in constitutive ISG expression (Fig. 4). IFN-I stimulation was marked by an increase in the number of ATAC-seq signals at ISG loci, which was strongly reduced in *Irf9-/-* macrophages (Fig. 4A, C). IFN-γ treatment also increased the accessibility at ISG loci, however loss of IRF9 did not markedly reduce the accessibility in this subset of genes (Fig. 4B, C).

In order to correlate ISGF3-dependent expression with accessibility changes in IFN-treated cells, we integrated RNA-seq and ATAC-seq data from wildtype (wt) and *Irf9-/-* macrophages. As previously published, basal expression of a subset of ISG dropped in *Irf9-/-* macrophages (Fig. 5A, colored dots, table EV2). These genes were further analyzed for corresponding changes in chromatin opening (red dots). *Oas1a, Oas2, Mx2* and *Mx1* were among the genes, that showed a drop in both expression and accessibility. *Irf1*, a target gene requiring STAT1 homodimer activity, was not affected by the loss of IRF9. Furthermore, we noted a high degree of overlap between genes whose expression was induced by IFN-I and IFN-γ (Fig. 5B,C; all colored dots, tables EV3, 4) and genes showing an increased accessibility by the respective treatment (red dots). For the majority of gene loci the type I IFN-dependent increase in chromatin accessibility required the presence of ISGF3 (472 blue dots out of 680 red dots in Fig. 5D, table EV 3). By comparison, a smaller fraction of loci with increased accessibility after IFN-γ stimulation was affected by the loss of IRF9 (153 blue dots out of 457 red dots in Fig. 5E, table EV4). The genes whose accessibility was affected by the absence of ISGF3 were scarce among the highly expressed genes. This, together with the fact that small changes of ATACseq signals are represented in the volcano plots of Fig. 5, explains why the gene group with increased accessibility contributed little to the profile of Fig. 4B. Among the ISGF3-dependent genes were typical core ISG from the *Mx* and *Oas* families (Fig. 5D-F). By contrast, *Irf1* accessibility at and upstream of the TSS was only weakly affected by IFN and not at all by the loss of IRF9 (Fig. 5G) ^28^. Taken together our data support the view that a subset of genes that are transcriptionally controlled by ISGF3 also exhibit ISGF3-dependent chromatin accessibility at their genomic loci in homeostatic as well as in IFN-induced conditions.

**Fig. 5:**
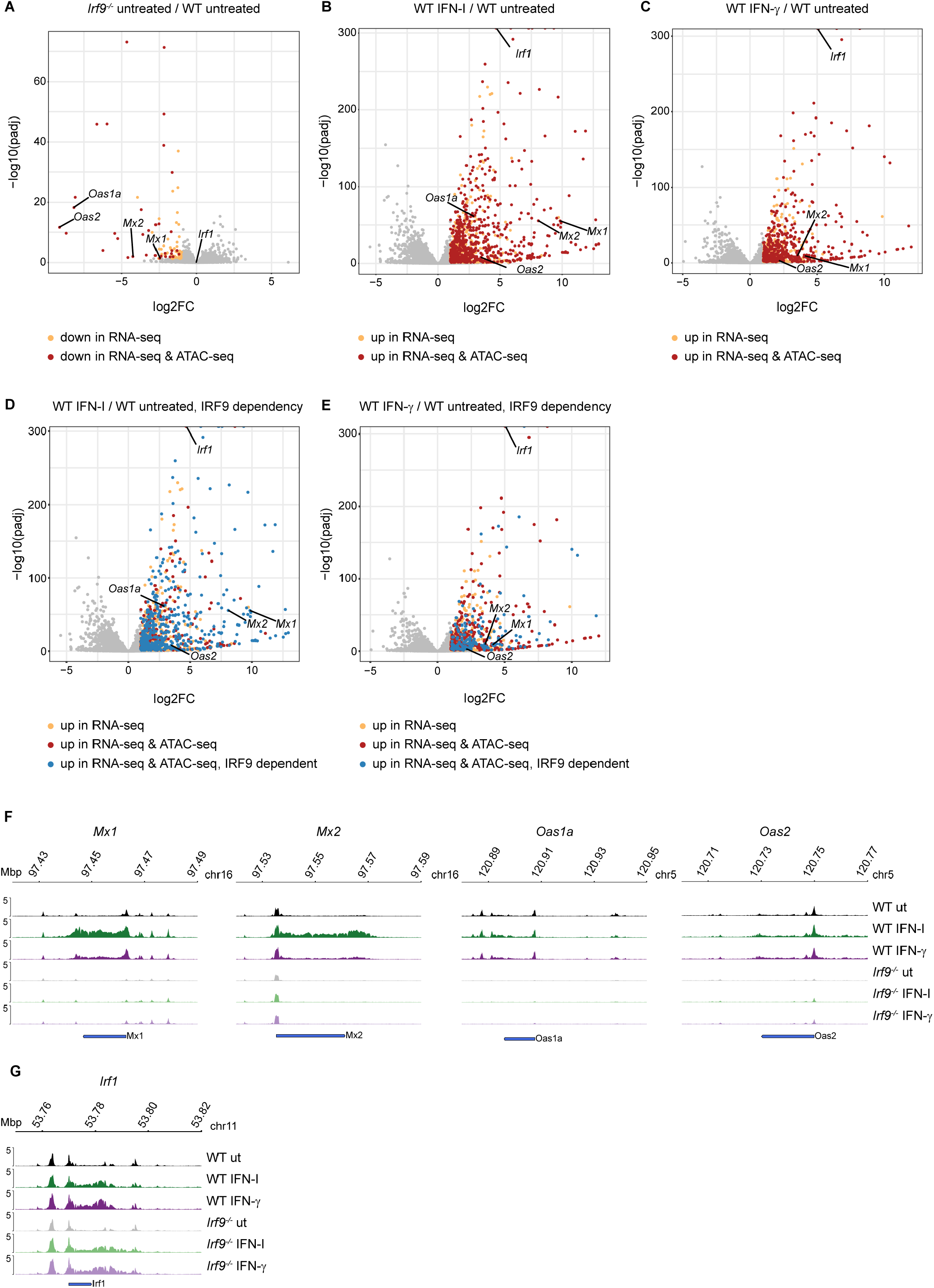
ISGF3 complex-dependent chromatin opening at ISG loci in homeostatic and interferon induced states. **A-F** Volcano plots of mRNA expression in BMDM in respective conditions. Each dot represents a gene. The log2-transformed fold change and -log10-transformed adjusted p-values for gene expression are shown in x and y axis respectively. **A** *Irf9*-/- vs wildtype BMDM in homeostatic condition: yellow dots represent genes significantly downregulated in *Irf9*-/-according to RNA-seq (log2FC ≤ 1, padj ≤ 0.05) and red dots represent genes which in addition showed significantly decreased accessibility at their respective loci according to ATAC-seq in *Irf9*-/- BMDM (log2FC ≤1, padj ≤ 0.05). **B** IFN-I (2h) stimulated vs untreated BMDM; **C** IFN-γ (2h) stimulated vs untreated BMDM. In the panels, yellow dots represent the genes which were significantly upregulated after either IFN treatment (log2FC ≥1, padj ≤0.05) according to RNA-seq. The red dots represent genes which in addition showed significant increase in chromatin accessibility (log2FC ≥1, padj ≤ 0.05) after corresponding IFN treatment at their genomic loci according to ATAC-seq data. **D, E** The blue dots represent genes which in addition, showed a significant decrease (log2FC ≥1, padj ≤ 0.05) in chromatin accessibility at their genomic loci in *Irf9*-/-BMDM in response to 2h of either IFN treatment (ATAC-seq log2FC ≥1, padj ≤ 0.05) **F, G** Genome browser tracks for ATAC-seq at respective gene loci in wildtype and *Irf9*-/- BMDM treated with IFN-I or IFN-γ for 2h (one representative replicate). Data in A-E are derived from three biological replicates.

### ISGF3 controls deposition of active histone marks and transcriptional activation at a subset of ISG

The changes in macrophage genome structure occurring in response to challenges such as an infection, ensure accessibility of critical regulatory DNA, thus allowing transcription factor binding and transcription of proinflammatory genes ^41^. An additional layer of gene control is formed by writers and erasers of histone marks.

We determined changes of activating or repressive histone marks with the aim of establishing whether both histone remodeling and modification at ISG promoters require ISGF3. To this end, we mined our IRF9 and RNA polymerase II (Pol II) ChIP-seq data as well as a publicly available H3K27ac ChIP-seq dataset ^28,43,44^. In addition, we performed site-directed ChIP using antibodies against H3K27ac, H3K27me3 and IRF9 in resting and IFN treated wt and *Irf9-/-* macrophages. IRF9 binding at *Mx1* and *Mx2* promoters is shown as a positive control (Fig. EV5A).

In line with previous observations ^28^, we noted different degrees of IRF9 pre-association with the promoters of the *Mx1, Mx2, Oas1a* and *Oas2* genes in resting cells (Fig. 6A, C, E, G). This correlates with reduced chromatin accessibility in resting *Irf9-/-* BMDM (Fig. 5F-I) and agrees with our recent finding showing that the STAT2-IRF9 complex controls basal ISG expression. The presence of H3K27ac was enriched at *Mx1, Mx2, Oas1a,* and *Oas2* gene promoters in response to both type I and type II IFN stimulation in wildtype cells and binding was significantly reduced in *Irf9-/-* BMDMs (Fig. 6B, D, F, H). In contrast, H3K27me3 was mostly unchanged in response to IFN-treatment, but all promoters except that of *Mx2* showed increased methylation in *Irf9-/-* macrophages. Hence, an IFN-independent, homeostatic ISGF3 activity is critically involved in removing repressive marks.

**Fig. 6:**
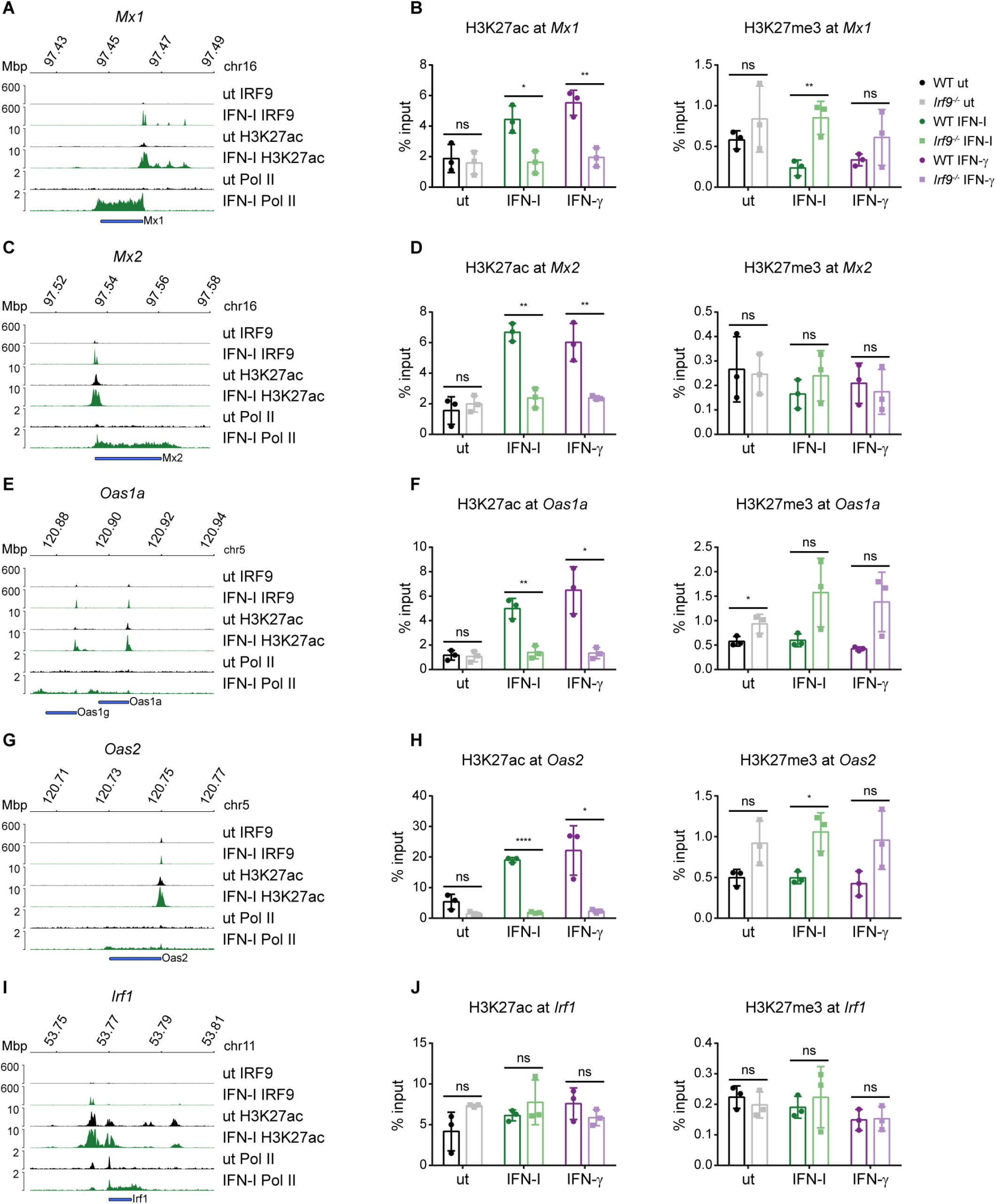
Histone modifications at ISG promoters correlates with the local chromatin accessibility. A, C, E, G, I Genome browser tracks representing published ChIP-seq datasets in BMDM with antibodies against the following proteins after IFN-I stimulation: IRF9 (1,5h), H3K27ac (1h) and Pol II (2h) showing promoter occupancy at indicated gene loci (one representative replicate). **B, D, F, H, J** Site directed ChIPs (*Mx1, Mx2, Oas1a, Oas2* and *Irf1*) were performed in bone marrow-derived macrophages isolated from wild-type and *Irf9*-/- BMDM, treated for 2h either with IFN-I or IFN-γ and processed for ChIP with antibodies targeting H3K27ac and H3K27me3 as indicated.

The pattern of H3K27ac deposition and Pol II loading was highly correlated to the promoter accessibility of the selected genes (Fig. 6A, C, E, G, I). Notably, at the *Irf1* gene, whose promoter is constitutively accessible independently of IFN-stimulation and the ISGF3 complex, the histone modifications did not change significantly after IFN treatment or upon IRF9 deficiency (Fig. 6I, J). Among the examined genes, this was also the only ISG with a preloaded Pol II.

Collectively, our findings demonstrate an ISGF3-complex dependent promoter activation in response to both IFN-I and IFN-γ, leading to increased chromatin accessibility, deposition of active histone marks and transcriptional activation at a subset of ISG.

## Discussion

3D chromatin structure is vital for both the organization and function of our genomes. Mitosis most likely represents the biggest challenge to the proper folding and positioning of chromatin in the nuclei of dividing cells, but 3D organization is of similar importance also for gene expression, i.e. the spatial arrangement of genes in active or inactive compartments and the controlled juxtaposition of regulatory elements in the process of transcriptional activation ^18^. Here, we investigated JAK-STAT signaling and its rapid control of nuclear ISG responses. We show that the structure of ISG loci undergoes similarly rapid changes: a reorganization of their loop formation and a movement or eviction of nucleosomes that increases the accessibility of the DNA containing regulatory elements. These changes are accompanied by the establishment of activating histone marks and, at most loci, the de novo recruitment of Pol II. Our studies are a starting point for future investigations into the precise mechanisms of chromatin reorganization at ISG loci.

The loop contact maps established in our Hi-C analyses show clustered ISG within the same TAD. This suggests that the chromatin loops at these loci may form and rearrange in accordance with the loop extrusion model, the assembly of cohesin/NIPBL/MAU2 complexes at binding sites for transcription factor CTCF and the subsequent ATP-dependent extrusion of DNA loops ^24,45–47^. The size of such loops is determined by the cohesin release factor WAPL^48^. WAPL repression by transcription factors may influence loop extrusion, as has been demonstrated for Pax 5 at the Ig heavy chain locus ^47^. The loop extrusion model thus provides an arena for further studies of the mechanisms by which IFN rearrange the chromatin of ISG loci. It is in agreement with a recent report about the importance of cohesin in the transcriptional response of macrophages to LPS ^50^.

Currently we can only speculate about mechanistic implications of IFN-induced or disrupted loops. With many ISG and their ISRE-containing promoters being organized in clusters, it is tempting to speculate that 3D rearrangements serve to bring the regulatory elements in close proximity. This may in turn create a high local concentration of ISGF3 complexes with the ability to exert their activity upon all genes united in the cluster. Some of the recorded intra-cluster loops are consistent with this idea. However, testing this hypothesis will involve the editing of ISRE sequences in their genomic context and to study the implications for transcriptional activation and the structure of the corresponding ISG locus.

Previous studies addressing chromatin remodeling at certain ISG loci may occur either before or after stimulation with IFN ^51–53^. Our ATAC-seq data now demonstrate on a genome-wide scale that IFN treatment opens chromatin at a large fraction of ISG promoters, but that some were rendered accessible independently of IFN. In addition, the results in IRF9-deficient cells clearly show that ISGF3 complexes are part of the machinery for chromatin remodeling at ISG. However, this is again not generally the case for all ISG. Thus, ISGF3 complexes may form complexes with and help to recruit chromatin remodelers as has been suggested ^54,55^. On the other hand, some genes manage to open their promoter chromatin via ISGF3-independent mechanisms both during homeostasis and after exposure to IFN. Generally speaking, ISG fall into the four categories: i) homeostatic accessibility, ISGF3-dependent, ii) homeostatic accessibility, ISGF3-independent, iii) IFN-induced accessibility, ISGF3-dependent and iv) IFN-induced accessibility, ISGF3-independent. We also noted that open chromatin at gene bodies frequently correlates with ongoing ISG transcription. This observation agrees with a report in virus-infected cells showing transcription-dependent chromatin rearrangements in influenza virus-infected cells ^56^.

Given the distinct activities of the two IFN types in the immune system, the degree of overlap between IFN-I and IFN-γ-induced transcriptomes is unexpected. To our surprise, similarities between the molecular mechanisms of transcriptional control by the two cytokines extended to all levels of regulation, from 3D loop rearrangement and promoter accessibility to histone modification and Pol II recruitment. In part, this may reflect the activity of the ISGF3 complex which shows the expected importance in gene regulation by IFN-I, but a larger than expected impact on IFN-γ-induced gene control. Compared to IFN-I signaling, STAT2 tyrosine phosphorylation downstream of the IFN-γ receptor is much less pronounced ^28,57^. In fact, the unphosphorylated fraction of STAT2 was shown to block nuclear entry of phosphorylated STAT1 through formation of hemi-phosphorylated heterodimers and thus to dampen the response to IFN-γ ^57^. In light of these findings it is surprising that most ISG responding to both IFN types show strong, IFN-γ-induced association with ISGF3 by ChIP-seq ^28^ and, consistently, a critical requirement for ISGF3 in producing open chromatin, depositing active histone marks and recruiting Pol II. One possible explanation for the discrepancy between little cytoplasmic STAT2 phosphorylation and the comparably large availability of nuclear ISGF3 for gene regulation could be trapping of nuclear complexes either by unspecific interaction with DNA, chromatin proteins, or the nuclear matrix. Trapping of ISGF3 may again increase its local concentration at specific locations of the genome.

Why then is the immunological activity of IFN-γ so different from that of IFN-I, particularly in macrophages? An important explanation may be provided by the rather small group of ISGF3-independent early response genes such as *Irf1. Irf1* is a first-tier gene encoding a transcription factor with a large contribution to the secondary IFN-γ regulome ^58,59^. This differs from IFN-I-induced genes which are largely independent of secondary IRF1 effects ^60^. Consistent with the notion that the *Irf1* gene is a primary response gene to IFN that is regulated independently of ISGF3, promoter chromatin opening, the deposition of activating marks and the recruitment of Pol II occurred under homeostatic conditions. This situation is reminiscent of primary, LPS-induced genes that are regulated at the level of transcription elongation ^61^. The group of secondary, IRF1-dependent IFN-γ target genes includes antimicrobial effector genes such as *Nos2* (*iNos*) and contributes to the macrophage-activating properties of IFN-γ ^59^. IRF8 is another STAT1 homodimer-dependent gene and important regulator of the IFN-γ-induced transcriptome in myeloid cells ^59,62^. It’s induction by IFN is much weaker than that of IRF1 but it plays an important role in the acquisition of transcriptional permissiveness at IFN-responsive enhancers ^44,62^. According to our data set the *Irf8* locus shows the same IFN/ISGF3 independence in chromatin accessibility as the *Irf1* locus. Thus, while there is clearly a large overlap and shared ISGF3-dependent regulome between IFN-I and IFN-γ that is responsible for common biological activities such as the antiviral state or regulation of inflammation, STAT1 homodimers, as drivers of IRF1 and IRF8 synthesis, are critical determinants for the role of IFN-γ as a macrophage-activating factor. While our study was focused on the immediate response to IFN-I and IFN-γ, delayed stages of the response to the cytokines are most likely characterized by larger differences between their induced transcriptomes, owing to a different deployment of transcription factors like IRF1 that determine secondary gene induction. Furthermore, IFN-γ reportedly promotes macrophage activation by selective repression of LPS-induced genes ^63^.Thus, the interplay and integration of the different levels of gene control downstream of the IFN-induced JAK-STAT pathways will remain an interesting topic for future studies.

## Material and Methods

### Mice, animal experiments

C57BL/6N mice (wildtype, *wt*) were purchased from Janvier Labs. *Irf9-/-* mice ^64^ were backcrossed for more than 10 generations on a C57BL/6N background. Mice were housed under identical conditions in a specific-pathogen-free (SPF) facility according to the Federation of European Laboratory Animal Science Association (FELASA) guidelines. Mice were bred under the approval of the institutional ethics and animal welfare committee of the University of Veterinary Medicine of Vienna and the national authority Federal Ministry Republic of Austria Education Science and Research section 8ff of the Animal Science and Experiments Act (Tierversuchsgesetz [TVG], BMWF-68.205/0068-WF/V/3b/2015 and GZ 2020-0.200.397). The study did not involve animal experiments as defined in the TVG and did not require ethical approval according to the local and national guidelines.

recommendations of the Federation of European Laboratory Animal Science Association and additionally monitored for being norovirus negative.

### Cell culture

Bone marrow-derived macrophages (BMDMs) were differentiated from bone marrow isolated from femurs and tibias of 8- to 12-week-old mice from female mice. Femur and tibia were flushed with Dulbecco’s modified Eagle’s medium (DMEM) (Sigma-Aldrich) and cells were differentiated in DMEM containing recombinant M-CSF (a kind gift from L. Ziegler-Heitbrock, Helmholtz Center, Munich, Germany). BMDMs were stimulated with 10 ng/ml murine IFN-γ (a kind gift from G. Adolf, Boehringer Ingelheim, Vienna) or 250 IU/mL of IFN-β (PBL Assay Science; Catalog # 12400-1).

### ChIP

1.5 × 10^7^ bone marrow derived macrophages were seeded on a 15 cm dish on day 7 of differentiation. The next day cells were stimulated for 1.5h either with 10 ng/ml murine IFN-γ (a kind gift from G. Adolf, Boehringer Ingelheim, Vienna) or 250 IU/mL of IFN-β (PBL Assay Science; Catalog # 12400-1). Cells were crosslinked for 10 minutes at room temperature in 1 % formaldehyde PBS (Thermo Fisher, Catalog # 28906). Cells were quenched with 0.125 M glycine for 10 min at RT. Cells were harvested and washed twice with ice cold PBS. Cells were centrifuged for 5 min at 1350 g at 4 °C. Pellets were snap frozen in liquid nitrogen and stored at 80 °C overnight. Frozen pellets were thawed on ice for 60 minutes. Pellets were resuspended in 5 mL LB1 (50 mM Hepes, 140 mM NaCl, 1 mM EDTA, 10 % glycerol, 0.5 % NP40, 0.25 % TritonX-100) by pipetting and rotated at 4 °C for 10 min. Samples were centrifuged for 5 minutes at 1350 x g at 4 °C. Pellets were resuspended in 5 mL LB2 (10 mM Tris, 200 mM NaCl, 1 mM EDTA, 0.5 mM EGTA) by pipetting and rotated at 4 °C for 10 min. Samples were centrifuged for 5 minutes at 1350 x g at 4 °C. Pellets were resuspended in 3 mL LB3 (10 mM Tris, 100 mM NaCl, 1 mM EDTA, 0.5 mM EGTA, 0.1 % deoxycholate, 0.5 % N-lauroylsarcosine). Samples were split into 2 x 1.5 mL in 15 mL polypropylene tubes suitable for the Bioruptor® Pico (Diagenode). BioRuptor Sonicator settings: power = high, “on” interval = 30 seconds, “off” interval = 45 seconds, 10 cycles. Sonicated samples were centrifuged for 10 minutes at 16000 x g at 4 °C to pellet cellular debris. Chromatin concentration was measured by NanoDrop and 25 µg of chromatin were used for each IP. 300 µl 10 % Triton X-100 were added to each 3 mL sonicated lysate. 25 µg of chromatin were stored at 4 °C which served later on as an input control. Antibody of interest was added to sonicated chromatin aliquot and mixed (anti-H3K27ac Cell Signaling, Catalog # 8173, 5μl; anti-H3K27me3 Active Motif, Catalog # 39155, 5ug; IRF9 6F1-H5 supernatant, 150 μl as previously described ^28^. All samples were filled up to 1ml with dilution buffer (16.5 mM Tris pH 8, 165 mM NaCl, 1.2 mM EDTA, 1 % Triton X-100, 0.1 % SDS, 0.1 mM PMSF, and complete EDTA-free protease inhibitor cocktail (Sigma Aldrich)). Samples were rotated at 4°C overnight.

50 µl of magnetic beads (Dynabeads protein G, Life technologies, 10003D) per sample were blocked overnight in dilution buffer containing 1 % BSA at 4 °C. The next day 50 µl of the beads were added to each sample and incubated at 4 °C while rotating. Afterwards the beads were washed 1x with RIPA buffer (50 mM Tris HCl pH 8, 150 mM NaCl, 1 % NP-40, 0.1 % SDS, 0.5 % sodium deoxycholate, 1 mM DTT), 2x high salt buffer (50 Mm Tris pH 8, 500 mM NaCl, 0.1 % SDS, 1 % NP-40), 2x LiCl buffer (50 mM Tris pH 8, 250 mM LiCl, 0.5 % sodium deoxycholate, 1 % NP-40) and TE buffer (10 mM Tris pH 8, 1 mM EDTA) for 10 minutes at 4 °C. The samples were eluted in freshly prepared elution buffer (2 % SDS, 100 mM NaHCO_3_, 10 mM DTT). The crosslink between proteins and DNA was reversed by adding 200 mM NaCl to each sample and incubation at 65 °C at 300 rpm for 12 hours. Proteinase K, 40 mM Tris pH 8 and 10 mM EDTA were added to each sample and incubated for 1 hour at 55 °C and 850 rpm. Each sample was transferred to a phase lock tube (5Prime), mixed 1:1 with phenol-chloroform-isoamylalcohol (PCI) and centrifuged for 5 minutes at 12000 × g. Supernatant was transferred and mixed with 800 µl 96 % Ethanol, 40 µl 3 M CH_3_COONa pH 5.3 and 1 µl Glycogen and stored for at overnight at -20 °C. Samples were centrifuged for 45 minutes at 4 °C and 16000 × g. Pellets were washed in ice cold 70 % ethanol and dried at 65 °C, before diluting the DNA in H_2_O.

Real-time qPCR were run on the Mastercycler (Eppendorf). Primers for ChIP PCR listed in table EV5.

### ATAC-seq

3 × 10^6^ BMDM derived from female mice, were seeded in 6-cm non-treated tissue culture plates on day 7 of differentiation. The next day, cells were stimulated for 2h either with 10 ng/ml murine IFN-γ (a kind gift from G. Adolf, Boehringer Ingelheim, Vienna) or 250 IU/mL of IFN-β (PBL Assay Science; Catalog # 12400-1).

Cells were gently washed with and scraped in PBS. A viability of >90% was determined with the cell viability analyzer and cell counter NucleoCounter® NC-3000™.

Cells were washed twice in 1 mL ice-cold PBS, by centrifuging cells at 1200 g for 5 minutes. Cells were resuspended in 1 mL cold nuclei isolation buffer (0.32 M sucrose; 3 mM CaCl2; 2 mM Mg Acetate; 0.1 mM EDTA; 10 mM Tris.HCl pH 8.0; 0.6% NP-40; 1mM DTT fresh) and placed on ice for 5 minutes. Cells were centrifuged at 700g for 5 min at 4°C. Supernatant was removed, 500 µL cold nuclei isolation buffer was added and samples were kept on ice for 3 min to further increase the number of nuclei isolated. Cells were centrifuged at 700g for 5 min at 4°C.

Supernatants were removed and pelleted nuclei were gently resuspended in 200 µL cold Nuclei Resuspension Buffer (NRB) (50 mM Tris.HCl pH 8.3; 40% Glycerol; 5 mM MgCl2; 0.1 mM EDTA). 50.000 nuclei (5µl) were resuspended in 7. 5µl H_2_O, 12.5 µl TD buffer and 5 µl Tn5 transposase from Illumina (#15027865).

Nuclei were incubated for 30 min at 37 °C, while mixing on a plate shaker (600 rpm). DNA was purified with Qiagen MinElute columns (#28006) according to the manufacturer’s instructions and eluted in 13µl of water. 12.5 µl purified DNA from the previous step were mixed with 5 µl 2.5 µM I7 index primer, 5 µl 2.5 uM I5 index primer (dual indexing), 2.5 µl Evagreen and 25 µl, 2x Q5 PCR Master Mix, (NEB # M0492L). An end point PCR was run as follows: 5 min 72 °C, 1 min 98 °C, 10sec 98 °C (5-7 cycles) 30sec 65 °C (5-7 cycles), 60sec 72 °C (NEB # E7645S).

The reaction was purified by adding 15 µl of SPRI beads prepared by the NGS facility (MBSpure beads) and incubated for 5 min at RT. The supernatant was transferred to a fresh well. 50 µl of MBSpure beads were added and mixed well. Incubated for 5 min at RT, magnetized, and removed supernatant. The beads were washed twice by adding 150 µl of 80 % EtOH. Beads were dried for 30 sec and DNA was eluted in 20 µl of water. The quality of the libraries was checked on a bioanalyzer to further determine the size distribution.

Libraries were sequenced on a NovaSeq6000 S2 PE50.

### Hi-C

Chromatin interaction libraries were generated using the Arima Genomics (https://arimagenomics.com/) Arima-Hi-C kit (P/N A510008).

2 × 10^6^ BMDM derived from female mice, were seeded in 6-cm non-treated tissue culture plates on day 7 of differentiation. The next day, cells were stimulated for 2h either with 10 ng/ml murine IFN-γ (a kind gift from G. Adolf, Boehringer Ingelheim, Vienna) or 250 IU/mL of IFN-β (PBL Assay Science; Catalog # 12400-1). Cells were gently washed with and scraped in PBS. A viability of >90% was determined with the cell viability analyzer and cell counter NucleoCounter® NC-3000™.

Cells were centrifuged at 500 g for 5 min and resuspended in 4.375 mL of 1xPBS at room temperature. 625 µl 16 % formaldehyde were added, mixed and incubated at RT for 10 min. 460 µl Stop Solution 1 was added, mixed well by inverting 10 times and incubated at RT for 5 min. Cells were placed on ice for 15 min and then pelleted by centrifugation (500 g, 5 min, 4 °C). Cells were resuspended in 2ml PBS and 1ml (1×10^6^ cells) were pelleted again (500 g, 5 min, 4 °C). The pellets were snap frozen in liquid nitrogen. Pellets were resuspended in 20 µl Lysis buffer and incubated at 4 °C for 15 min. 24 µl conditioning solution was added, gently mixed and incubated at 62 °C for 10 min. 20 µl of Stop solution 2 were added, mixed and incubated for 15 min at 37 °C. A mix of 7 µl Buffer A, 1 µl enzyme A1 and 4 µl of enzyme A2 were added to the sample and incubated as follows: 37 °C for 60 min, 65 °C 20 min, 25 °C 10 min. 12 µl of Buffer B and 4 µl of enzyme B were added to the mix and incubated for 45 min at RT. Further 70 µl Buffer C and 12 µl enzyme C were added, mixed gently and incubated for 15 minutes at RT. 10,5 µl Buffer D and 25 µl enzyme D were mixed. 20 µl Buffer E were added and the whole mix was added to the sample and incubated as follows: 55 °C for 30 min, 68 °C for 90 min, 4 °C hold. 100 µl of MBS pure beads were added and incubated at RT for 5 min. The supernatants were removed from the magnetized solution. 200 µl 80 % EtOH were added while the plate was still on the magnet. EtOH was removed, beads were resuspended in 50 µl elution buffer, and incubated at RT for 5 min. The eluate was transferred to a new tube and the DNA concentration was determined by Qubit. 5 µg DNA of each sample were fragmented by using the Covaris S2 and the following program: Intensity - 5, Duty Cycle - 10%, Cycles per burst – 200, Treatment Time - 50 sec, Temperature – 7 °C. 50 µl of MBS pure beads were added to the 100 µl of sheared solution and incubated for 5 min at RT. Supernatants from magnetized solutions were transferred to a fresh well. 50 µl of beads were added and incubated for 5 min at RT. Supernatant was removed from magnetized solution and 200 µl 80 % EtOH were added, while the plate was still on magnet. EtOH was removed and beads were resuspended in 100 µl elution buffer. DNA amount was quantified by Qubit and proceeded to Biotin enrichment protocol and library preparation. 300ng of DNA were used for the library preparation using the Ultra II kit (NEB) for adaptation. 100 µl of enrichment beads were added and incubated at RT for 15 min. Supernatant was discarded, beads were resuspended in 180 µl wash buffer and incubated at 55 °C for 2 min. The washing step was performed again. Beads were resuspended in 100 µl elution buffer and the supernatant was again removed. Beads were resuspended in 50 µl elution buffer, 7 µl End prep buffer and 3 µl end prep enzyme were added and the mix was incubated 30 min at 20 °C and 30 min at 65 °C. 30 µl Ligation master mix, 2.5 µl Adapter and 1 µl ligation enhancer were added and incubated at RT for 15 min. The beads were magnetized and the supernatant was discarded. Beads were washed twice in 180 µl Wash buffer, while incubated at 55 °C for 2 min. Beads were resuspended in 100 µl elution buffer at RT. Supernatants were removed and beads were resuspended in 12 µl of elution buffer. 3 µl of USER enzyme, 10 µl i5/i7 index primer mix (2.5 µM), 2.5 µl Evagreen and 25 µl Q5 PCR mix from NEB ultra II Kit (E7645S) were added and the PCR was run as follows: 37 °C 15 min, 98 °C 30 sec, 98 °C 10 sec, 65 °C 75 sec (6 cycles endpoint PCR). Libraries were purified with 1x MBS pure beads. Libraries were sequenced on a NovaSeq6000 S1 PE50.

### Hi-C data processing

Hi-C data was processed using the Hi-C pipeline from the nf-core framework (nf-core/Hi-C v1.1.0; https://zenodo.org/record/2669513#.YA8UvWRKjZk; ^65^). Reads were aligned against the Illumina iGenome *Mus musculus* GRCm38 reference genome. Restriction sites used were ^GATC, G^ANTC. Ligation sites used were GATCGATC, GANTGATC, GANTANTC, GATCANTC.

### HiCreppy

Sample similarity for Fig. EV1A was assessed using hicreppy (v0.0.6) (https://github.com/cmdoret/hicreppy), a reimplementation of HiCRep (doi: 10.1101/gr.220640.117), for Hi-C maps with 10 kb bins and a maximum scanning distance of 1 Mb. The algorithm is computing a stratum-adjusted correlation coefficient, which was used for evaluating reproducibility across replicates and conditions.

### PCA/Compartment analysis

The first three eigenvectors of Hi-C contact maps were computed with *cooltools.eigdecomp* (cooltools, v0.3.0) using bedgraph tracks from published RNA-seq data ^28^ to phase and rank them according to their correlation. The eigenvector showing the highest correlation with the RNA-seq data was considered the one representing A and B compartments. Vectors were processed independently for each chromosome. In order to measure the compartmentalization strength, we created compartmentalization plots (“saddle plot”) with Hi-C maps at 80 kb resolution using cooltools, as described in their manual (https://cooltools.readthedocs.io/en/latest/index.html).

### Ratio plots

Ratio plots, comparing interaction strengths between two conditions (Fig. 1B, D) were generated using Serpentine ^66^, a flexible 2D binning method improving especially the comparison of poorly covered and noisy regions. It is operating on raw data and is only performing binning where necessary on noisy regions, while at the same time preserving the resolution of highly covered regions.

### Pattern detection

Pattern detection in the Hi-C contact maps was performed using Chromosight (v1.3.3) ^35^. Chromosight is sliding pattern templates over contact maps and, for each position, computing Pearson correlation coefficients between the template and the contact map.

In order to make the correlation scores for called patterns comparable between replicates and conditions, a fusion map was created containing the contacts of all replicates across conditions with *cooler* (v0.8.10) ^67^. A first round of pattern detection was performed on this fusion map with *chromosight detect* on contact maps with bin sizes of 5 kb and 10 kb at a *maximum scanning distance* of 10 Mb. The pattern used was *loops_small.* The positions of detected patterns were passed to *chromosight quantify* for quantification of Pearson correlation scores in individual contact maps for each condition. Therefore, contact maps were subsampled to the number of contacts in the smallest matrix to ensure equal coverage across conditions. For Fig. 2A-F, Fig. 3A, B, Fig. EV2A, B and Fig. EV3A-C detected patterns were filtered for Pearson scores > 0.35.

### ATAC-seq data processing

ATAC-seq data was processed using the ATAC-seq pipeline from the nf-core framework(<u>nf-core/atacseq</u>v1.2.1, https://zenodo.org/record/3965985#.YA8a8GRKjZk). Reads were aligned against the Illumina iGenome *Mus musculus* GRCm38 reference genome. Downstream analysis of ATAC-seq data was performed using MACS2 narrow-peak calling and differential chromatin accessibility analysis (included in nf-core/atacseq pipeline).

### ATAC-seq heatmaps

For the generation of summary profile plots and heatmaps, density information (bigwig) for gene regions and surrounding regions (2 kbp upstream of TSS and 1 kbp downstream of TES) was plotted using *deeptools* (v3.4.3) (https://zenodo.org/record/3965985#.YA8XPGRKjZk). For defining clusters of ISG, we filtered the top 200 significantly (p-value ≤ 0.05) upregulated genes (sorted by log2FC) according to RNA-seq analysis. The second cluster refers to the respective remaining genes in the genome. The *computeMatrix* command was used in the *scale-regions* mode with the option *missingDataAsZero*. Subsequently, we used *plotHeatmap* for the generation of heatmaps and summary profiles.

### ATAC-seq integration with RNA-seq

For integrating ATAC-seq and RNA-seq we used the respective differential analyses from DESeq2. For ATAC-seq, each called interval (available output from the nf-core pipeline) was annotated to a gene by using the HOMER script *annotatePeaks.pl*. ^68^ Thereby peaks were annotated to genes and could be compared with the RNA-seq analysis.

### ChIP-seq data processing

Published ChIP-seq data ^28,43,44^ was re-analysed using the ChIP-seq pipeline from the nf-core framework (nf-core/chipseq v1.1.0) (https://zenodo.org/record/3529400#.YA8YGmRKjZk). Reads were aligned against the Illumina iGenome *Mus musculus* GRCm38 reference genome.

### Statistical analysis

Site directed ChIP data represent the mean values with standard deviation (SD). Statistical significance was calculated using two-tailed unpaired t-test. All statistical analysis was performed using GraphPad Prism (Graphpad) software. Asterisks denote statistical significance as follows: ns: p > 0.05, *: p ≤ 0.05, **: p ≤ 0.01, ***: p ≤ 0.001. Each dot represents one biological replicate.

### Data availability

Publicly available raw data reported in this paper are available under accession numbers GEO: GSE115435 (IRF9 ChIP), ArrayExpress: E-MTAB-2972 (Pol II),GEO: GSE56123 (H3K27Ac). Raw data generated for this publicationare available under SRA BioProject accession PRJNA694816.

## Acknowledgements

We thank Carlo Pecoraro and Physalia Courses (www.physalia-courses.org) for all arranged NGS data analysis training. We are especially grateful to Federico Comoglio and Cyril Matthey-Doret for sharing their expertise and enthusiasm for data visualization, ATAC-Seq and Hi-C-analysis with us. We thank Manuela Baccarini for critical comments to the manuscript and R. Stocsits for suggestions on Hi-C analysis. Solexa sequencing was performed by the VBCF NGS Unit (www.vbcf.ac.at). Funding was provided by the Austrian Science Fund (FWF) through project P25186-B22 to TD and SFB F6101, F6103 and F6106 to TD and MM.

## Author contribution

E.P. and T.D. conceived and developed the outline of this research and supervised the project. E.P. carried out the Hi-C and ATAC-seq experiments and contributed to data analysis. S.G. analyzed the Hi-C data, ATAC-seq data, reanalyzed ChIP-seq data and prepared figures. A.G. contributed to data analysis, ATAC-seq experiments and figure preparation. L.B. carried out the ChIP experiments and their statistical analysis. A.V. assisted in Hi-C and ATAC-seq sample preparation and protocol optimization. M.N. carried out the pre-processing of the Hi-C dataset. A.S. and M.M. provided intellectual input and expertise. E.P., S.G., A.G. and T.D. helped shape the research and analysis and wrote the paper. All authors read and commented on the paper.

## Figure legends

**Fig. EV1**

**A** Heatmap of sample similarity, as calculated with HiCRep (max. scanning distance 10Mbp, 10kbp bins). **B-D** Density plots of RNA-seq read coverage versus compartment score for each genomic bin (40kbp) of **B** untreated, **C** IFN-I treated (2h), **D** IFN-γ treated (2h) primary murine bone marrow-derived macrophages (BMDM). **E-G** Compartmentalization saddle plots showing the enrichment of interactions as a function of the compartment vector in **E** untreated, **F** IFN-I (2h) and **G** IFN-γ (2h) treated primary murine bone marrow-derived macrophages (kbp bins).

**Fig. EV2**

**A, B** Hi-C contact maps of untreated and IFN-I treated BMDM. Visualization of pattern detection with chromosight (loops between two loci are indicated as arcs). Color coding of arcs corresponds to Pearson correlation scores. Loops with a score > 0.35 are shown (maximum size/ylim = 500 kbp). Only ISG of the corresponding gene clusters are visualized. Single replicates are plotted next to each other. **c** HiC contact maps (merge of 2 replicates per condition, log1p scale) of untreated and IFN-I treated (2h) BMDM. Visualization of pattern detection with chromosight (loops between two loci are indicated as arcs). Color coding of arcs corresponds to Pearson correlation scores. Loops with a score > 0.35 are shown (maximum size/ylim = 500 kbp). Lower panel, IRF9 ChIP-seq tracks after IFN-I (1.5h) stimulation and gene annotations are shown to visualize ISGF3 binding sites. Only ISG of the corresponding gene clusters are visualized.

**Fig. EV3**

**A-D** Hi-C contact maps (merge of 2 replicates per condition, log1p scale) of untreated and IFN-γ (2h) treated BMDM. Visualization of pattern detection with chromosight (loops between two loci are indicated as arcs). Color coding of arcs corresponds to Pearson correlation scores. Loops with a score > 0.35 are shown (maximum size/ylim = 500 kbp). Lower panel, STAT1 ChIP-seq tracks (1.5h IFN-I treatment) and gene annotations. Only ISG of the corresponding gene clusters are visualized.

**Fig. EV4**

**A** Heatmap of sample similarity, generated from clustering by Euclidean distances between DESeq2 rlog values for each sample. **B, C** Heatmaps of chromatin accessibility in regions of the top 200 genes (log2FC) upregulated after IFN-I (2h) and IFN-γ (2h) treatment respectively according to RNA-seq and the remaining genome. Heatmaps are shown for the regions of the genes, including 2 kbp upstream of the TSS and 1kbp downstream of the TES as indicated below the plots. Data are derived from three biological replicates.

**Fig. EV5**

**A** Site directed ChIP at the *Mx1, Mx2* gene promoters was performed with anti-IRF9 mAb using bone marrow-derived macrophages isolated from wild-type and *Irf*9-/- BMDM, treated for 2h either with IFN-I or IFN-γ.

## References

1. MacMicking, J. D. Interferon-inducible effector mechanisms in cell-autonomous immunity. Nature Reviews Immunology (2012) doi:10.1038/nri3210

2. McNab, F., Mayer-Barber, K., Sher, A., Wack, A. & O’Garra, A. Type I interferons in infectious disease. Nature Reviews Immunology (2015) doi:10.1038/nri3787

3. Schneider, W. M., Chevillotte, M. D. & Rice, C. M. Interferon-stimulated genes: A complex web of host defenses. Annual Review of Immunology (2014) doi:10.1146/annurev-immunol-032713-120231

4. Levy, D. E. & Darnell, J. E. STATs: Transcriptional control and biological impact. Nature Reviews Molecular Cell Biology (2002) doi:10.1038/nrm909

5. Darnell, J. E., Kerr, I. M. & Stark, G. R. Jak-STAT pathways and transcriptional activation in response to IFNs and other extracellular signaling proteins. Science (80-.). (1994) doi:10.1126/science.8197455

6. Sancho-Shimizu, V., Perez De Diego, R., Jouanguy, E., Zhang, S. Y. & Casanova, J. L. Inborn errors of anti-viral interferon immunity in humans. Current Opinion in Virology (2011) doi:10.1016/j.coviro.2011.10.016

7. Crow, Y. J. & Manel, N. Aicardi-Goutières syndrome and the type I interferonopathies. Nature Reviews Immunology (2015) doi:10.1038/nri3850

8. Stark, G. R. & Darnell, J. E. The JAK-STAT Pathway at Twenty. Immunity (2012) doi:10.1016/j.immuni.2012.03.013

9. Au-Yeung, N., Mandhana, R. & Horvath, C. M. Transcriptional regulation by STAT1 and STAT2 in the interferon JAK-STAT pathway. JAK-STAT (2013) doi:10.4161/jkst.23931

10. Kessler, D. S., Veals, S. A., Fu, X. Y. & Levy, D. E. Interferon-α regulates nuclear translocation and DNA-binding affinity of ISGF3, a multimeric transcriptional activator. Genes Dev. (1990) doi:10.1101/gad.4.10.1753

11. Schindler, C., Levy, D. E. & Decker, T. JAK-STAT signaling: From interferons to cytokines. Journal of Biological Chemistry (2007) doi:10.1074/jbc.R700016200

12. Ivashkiv, L. B. & Donlin, L. T. Regulation of type i interferon responses. Nature Reviews Immunology (2014) doi:10.1038/nri3581

13. Decker, T., Kovarik, P. & Meinke, A. GAS elements: A few nucleotides with a major impact on cytokine-induced gene expression. J. Interf. Cytokine Res. (1997) doi:10.1089/jir.1997.17.121

14. Maniatis, T., Goodbourn, S. & Fischer, J. A. Regulation of inducible and tissue-specific gene expression. Science (80-.). (1987) doi:10.1126/science.3296191

15. Smale, S. T. & Kadonaga, J. T. The RNA polymerase II core promoter. Annual Review of Biochemistry (2003) doi:10.1146/annurev.biochem.72.121801.161520

16. Schoenfelder, S. & Fraser, P. Long-range enhancer–promoter contacts in gene expression control. Nature Reviews Genetics (2019) doi:10.1038/s41576-019-0128-0

17. Bonev, B. & Cavalli, G. Organization and function of the 3D genome. Nature Reviews Genetics (2016) doi:10.1038/nrg.2016.112

18. Dekker, J. & Mirny, L. The 3D Genome as Moderator of Chromosomal Communication. Cell (2016) doi:10.1016/j.cell.2016.02.007

19. Lieberman-Aiden, E. et al. Comprehensive mapping of long-range interactions reveals folding principles of the human genome. Science (80-.). (2009) doi:10.1126/science.1181369

20. Rowley, M. J. & Corces, V. G. Organizational principles of 3D genome architecture. Nat. Rev. Genet. (2018) doi:10.1038/s41576-018-0060-8

21. Rao, S. S. P. et al. Cohesin Loss Eliminates All Loop Domains. Cell (2017) doi:10.1016/j.cell.2017.09.026

22. Schwarzer, W. et al. Two independent modes of chromatin organization revealed by cohesin removal. Nature (2017) doi:10.1038/nature24281

23. Buenrostro, J. D., Wu, B., Chang, H. Y. & Greenleaf, W. J. ATAC-seq: A method for assaying chromatin accessibility genome-wide. Curr. Protoc. Mol. Biol. (2015) doi:10.1002/0471142727.mb2129s109

24. Wutz, G. et al. Topologically associating domains and chromatin loops depend on cohesin and are regulated by CTCF, WAPL, and PDS5 proteins. EMBO J. (2017) doi:10.15252/embj.201798004

25. Dowen, J. M. et al. Control of cell identity genes occurs in insulated neighborhoods in mammalian chromosomes. Cell (2014) doi:10.1016/j.cell.2014.09.030

26. Symmons, O. et al. The Shh Topological Domain Facilitates the Action of Remote Enhancers by Reducing the Effects of Genomic Distances. Dev. Cell (2016) doi:10.1016/j.devcel.2016.10.015

27. Dixon, J. R. et al. Topological domains in mammalian genomes identified by analysis of chromatin interactions. Nature (2012) doi:10.1038/nature11082

28. Platanitis, E. et al. A molecular switch from STAT2-IRF9 to ISGF3 underlies interferon-induced gene transcription. Nat. Commun. (2019) doi:10.1038/s41467-019-10970-y

29. Majoros, A. et al. Canonical and non-canonical aspects of JAK-STAT signaling: Lessons from interferons for cytokine responses. Frontiers in Immunology (2017) doi:10.3389/fimmu.2017.00029

30. Yang, T. et al. HiCRep: assessing the reproducibility of Hi-C data using a stratum-adjusted correlation coefficient. Genome Res. (2017) doi:10.1101/gr.220640.117

31. Gough, D. J., Messina, N. L., Clarke, C. J. P., Johnstone, R. W. & Levy, D. E. Constitutive Type I Interferon Modulates Homeostatic Balance through Tonic Signaling. Immunity (2012) doi:10.1016/j.immuni.2012.01.011

32. Comoglio, F. et al. Thrombopoietin signaling to chromatin elicits rapid and pervasive epigenome remodeling within poised chromatin architectures. Genome Res. (2018) doi:10.1101/gr.227272.117

33. Jin, F. et al. A high-resolution map of the three-dimensional chromatin interactome in human cells. Nature (2013) doi:10.1038/nature12644

34. Le Dily, F. L. et al. Distinct structural transitions of chromatin topological domains correlate with coordinated hormone-induced gene regulation. Genes Dev. (2014) doi:10.1101/gad.241422.114

35. Matthey-Doret, C. et al. Computer vision for pattern detection in chromosome contact maps. Nat. Commun. (2020) doi:10.1038/s41467-020-19562-7

36. Rein, T., Müller, M. & Zorbas, H. In vivo footprinting of the IRF-1 promoter: Inducible occupation of a GAS element next to a persistent structural alteration of the DNA. Nucleic Acids Res. (1994) doi:10.1093/nar/22.15.3033

37. Pine, R., Canova, A. & Schindler, C. Tyrosine phosphorylated p91 binds to a single element in the ISGF2/IRF-1 promoter to mediate induction by IFNα and IFNγ, and is likely to autoregulate the p91 gene. EMBO J. (1994) doi:10.1002/j.1460-2075.1994.tb06245.x

38. Kanno, Y. et al. The genomic structure of the murine ICSBP gene reveals the presence of the gamma interferon-responsive element, to which an ISGF3 alpha subunit (or similar) molecule binds. Mol. Cell. Biol. 13, 3951–3963 (1993).

39. Parrini, M. et al. The C-Terminal Transactivation Domain of STAT1 Has a Gene-Specific Role in Transactivation and Cofactor Recruitment. Front. Immunol. 9, 2879 (2018).

40. Rauch, I. et al. Noncanonical Effects of IRF9 in Intestinal Inflammation: More than Type I and Type III Interferons. Mol. Cell. Biol. (2015) doi:10.1128/mcb.01498-14

41. Ostuni, R. et al. Latent enhancers activated by stimulation in differentiated cells. Cell (2013) doi:10.1016/j.cell.2012.12.018

42. Calderon, D. et al. Landscape of stimulation-responsive chromatin across diverse human immune cells. Nat. Genet. (2019) doi:10.1038/s41588-019-0505-9

43. Wienerroither, S. et al. Cooperative Transcriptional Activation of Antimicrobial Genes by STAT and NF-κB Pathways by Concerted Recruitment of the Mediator Complex. Cell Rep. (2015) doi:10.1016/j.celrep.2015.06.021

44. Mancino, A. et al. A dual cis-regulatory code links IRF8 to constitutive and inducible gene expression in macrophages. Genes Dev. (2015) doi:10.1101/gad.257592.114

45. Wutz, G. et al. ESCO1 and CTCF enable formation of long chromatin loops by protecting cohesinstag1 from WAPL. Elife (2020) doi:10.7554/eLife.52091

46. Davidson, I. F. et al. DNA loop extrusion by human cohesin. Science (80-.). (2019) doi:10.1126/science.aaz3418

47. Kim, Y., Shi, Z., Zhang, H., Finkelstein, I. J. & Yu, H. Human cohesin compacts DNA by loop extrusion. Science (80-.). (2019) doi:10.1126/science.aaz4475

48. Haarhuis, J. H. I. et al. The Cohesin Release Factor WAPL Restricts Chromatin Loop Extension. Cell (2017) doi:10.1016/j.cell.2017.04.013

49. Hill, L. et al. Wapl repression by Pax5 promotes V gene recombination by Igh loop extrusion. Nature (2020) doi:10.1038/s41586-020-2454-y

50. Cuartero, S. et al. Control of inducible gene expression links cohesin to hematopoietic progenitor self-renewal and differentiation. Nat. Immunol. (2018) doi:10.1038/s41590-018-0184-1

51. Ni, Z. et al. Apical role for BRG1 in cytokine-induced promoter assembly. Proc. Natl. Acad. Sci. U. S. A. (2005) doi:10.1073/pnas.0503070102

52. Mostafavi, S. et al. Parsing the Interferon Transcriptional Network and Its Disease Associations. Cell (2016) doi:10.1016/j.cell.2015.12.032

53. Cheng, Q. et al. Sequential conditioning-stimulation reveals distinct gene- and stimulus-specific effects of Type I and II IFN on human macrophage functions. Sci. Rep. (2019) doi:10.1038/s41598-019-40503-y

54. Gnatovskiy, L., Mita, P. & Levy, D. E. The Human RVB Complex Is Required for Efficient Transcription of Type I Interferon-Stimulated Genes. Mol. Cell. Biol. (2013) doi:10.1128/mcb.01562-12

55. Au-Yeung, N. & Horvath, C. M. Histone H2A.Z Suppression of Interferon-Stimulated Transcription and Antiviral Immunity Is Modulated by GCN5 and BRD2. iScience (2018) doi:10.1016/j.isci.2018.07.013

56. Heinz, S. et al. Transcription Elongation Can Affect Genome 3D Structure. Cell (2018) doi:10.1016/j.cell.2018.07.047

57. Ho, J. et al. STAT2 Is a Pervasive Cytokine Regulator due to Its Inhibition of STAT1 in Multiple Signaling Pathways. PLoS Biol. (2016) doi:10.1371/journal.pbio.2000117

58. Ramsauer, K. et al. Distinct modes of action applied by transcription factors STAT1 and IRF1 to initiate transcription of the IFN-γ-inducible gbp2 gene. Proc. Natl. Acad. Sci. U. S. A. (2007) doi:10.1073/pnas.0610944104

59. Langlais, D., Barreiro, L. B. & Gros, P. The macrophage IRF8/IRF1 regulome is required for protection against infections and is associated with chronic inflammation. J. Exp. Med. (2016) doi:10.1084/jem.20151764

60. Forero, A. et al. Differential Activation of the Transcription Factor IRF1 Underlies the Distinct Immune Responses Elicited by Type I and Type III Interferons. Immunity (2019) doi:10.1016/j.immuni.2019.07.007

61. Adelman, K. et al. Immediate mediators of the inflammatory response are poised for gene activation through RNA polymerase II stalling. Proc. Natl. Acad. Sci. U. S. A. (2009) doi:10.1073/pnas.0910177106

62. Platanitis, E. & Decker, T. Regulatory networks involving STATs, IRFs, and NFκB in inflammation. Frontiers in Immunology (2018) doi:10.3389/fimmu.2018.02542

63. Kang, K. et al. IFN-γ selectively suppresses a subset of TLR4-activated genes and enhancers to potentiate macrophage activation. Nat. Commun. 10, 1–14 (2019).

64. Kimura, T. et al. Essential and non-redundant roles of p48 (ISGF3$γ$) and IRF-1 in both type I and type II interferon responses, as revealed by gene targeting studies. Genes to Cells (1996) doi:10.1046/j.1365-2443.1996.08008.x

65. Ewels, P. A. et al. The nf-core framework for community-curated bioinformatics pipelines. Nature Biotechnology (2020) doi:10.1038/s41587-020-0439-x

66. Baudry, L., Millot, G. A., Thierry, A., Koszul, R. & Scolari, V. F. Serpentine: a flexible 2D binning method for differential Hi-C analysis. Bioinformatics 36, 3645–3651 (2020).

67. Abdennur, N. & Mirny, L. A. Cooler: Scalable storage for Hi-C data and other genomically labeled arrays. Bioinformatics (2020) doi:10.1093/bioinformatics/btz540

68. Heinz, S. et al. Simple Combinations of Lineage-Determining Transcription Factors Prime cis-Regulatory Elements Required for Macrophage and B Cell Identities. Mol. Cell (2010) doi:10.1016/j.molcel.2010.05.004

